# Role of GLP1-receptor signaling and α-β communication in functional β-cell heterogeneity

**DOI:** 10.1101/2025.04.25.650736

**Authors:** Nirmala V. Balasenthilkumaran, Lidija Križančić Bombek, Shane Browne, David Ramirez, Maja Duh, Maša Skelin Klemen, Eva Paradiž Leitgeb, Jasmina Kerčmar, Jan Kopecky, Yaowen Zhang, Maria Skjott Hansen, Yidi Shao, Annanya Sethiya, Adam Takaoglu, Divya Prabhu, Aining Fan, Shravani Vitalapuram, Christian M. Cohrs, Stephan Speier, Jurij Dolenšek, Rahaf Shishani, Anna Jackson, Bethany P. Cummings, Marko Gosak, Andraž Stožer, Vira Kravets

**Author notes:** corresponding author: Email ID. equal contributions (Nirmala V. Balasenthilkumaran, Lidija Križančić Bombek).

## Abstract

While islet β-cells were first viewed as a singular functional entity, since the 1970s findings reveal that individual β-cells differ in their insulin secretion. More recently distinct functional subpopulations based on differential calcium dynamics have been demonstrated to drive islet function. Here, we investigate how paracrine signaling, specifically glucagon-like peptide receptor (GLP-1R) signaling shapes functional β-cell heterogeneity. To address this, we utilized confocal imaging of calcium responses of in-vitro islets from MIP-CreER-GCaMP6s mice, C57BL/6 mice, and β-cell-specific humanized GLP-1R Cre-inducible KO mice. Responses in isolated islets vs pancreatic slices both before and after GLP-1R antagonist (exendin-9) or agonist (exendin-4) treatment were compared. We employ consecutive glucose stimulations in the same islet, where first stimulation serves as a control for the second stimulation. Function of the β-cells was significantly enhanced in the 2^nd^ glucose stimulation, reflecting previously reported time-dependent glucose potentiation phenomenon. GLP-1R antagonism delayed response time, decreased peak amplitude, active time and oscillation frequency, and disrupted the temporal consistency of hub-cells and 1^st^ responder β-cells, relative to the 1^st^ stimulation. Opposite effects were observed with the GLP-1R agonist. The changes in Ca^2+^ frequency were more pronounced in α-neighboring β-cells, suggesting local GLP-1R mediated α-β-cell communication. GLP-1R KO elicited deleterious effects, which, aside from the network connectivity, mirrored those of the ex-9. Together, these findings underscore the importance of GLP-1R-mediated α- and β-cell communication in heterogeneous β-cell function.

**Article Highlights:**

- Role of GLP-1R mediated α-β cell communication in functional β-cell heterogeneity was unclear.
- Time of response to glucose, 1^st^ phase amplitude, speed of signal propagation, active time, functional connectivity, and consistency of 1^st^ responder and hub cell subpopulations are enhanced by GLP-1R signaling.
- GLP-1R signaling creates local α-cell-associated functional neighborhoods of β-cells and maintains network organization.
- Islet-local GLP-1R communication in absence of exogenous GLP-1 is sufficient for significant control of β -cell function.

## Introduction

Insulin-producing β-cells exhibit different transcriptomic profiles, protein expression, and gap junction coupling (1–5). These variations result in a heterogeneous response to glucose stimulation, leading to the existence of distinct subpopulations of β-cells within the islet. We and others have previously shown the existence, temporal and spatial stability of unique functional β-cell subpopulations within the islet (6–8). Although recent studies have identified markers that could separate β-cell subpopulations, these markers are not always linked to β- cell function, leaving the mechanisms shaping these β-cell subpopulations poorly understood (5; 9; 10).

The islets also contain glucagon-secreting α-cells and somatostatin-secreting δ-cells, which interact through paracrine signaling (11; 12). Glucagon secreted by α-cells is sensed by glucagon and GLP-1 receptors on β-cells, enhancing glucose-stimulated cyclic AMP, calcium ([Ca^2+^]) influx, and insulin secretion (13; 14). We have previously shown that immune cells preferentially interact with α-cell-rich regions of an islet in pre-T1D, and α-cells have also been reported to have higher levels of HLA-I during early T1D, illustrating the importance of α-cells both in health and diabetes (15; 16). Understanding the extent to which α-cells influence β-cell function and heterogeneity in healthy conditions aids better understanding of diabetes pathophysiology.

Glucose stimulation increases intracellular ATP in β-cells, leading to membrane depolarization, [Ca^2+^] influx, and biphasic insulin release consisting of a transitional 1^st^ phase followed by an oscillatory 2^nd^ phase (17–20). While these quantitative functional parameters for these phases are well characterized (6; 7; 21; 22), the extent to which they are regulated by intra-islet GLP-1R signaling remains unclear. Here, we investigate the role of GLP-1R signaling in modulating β-cell’s 1^st^ and 2^nd^ phase response to glucose, and in shaping of the functional β-cell subpopulations.

In previous studies, the manipulation of α-β signaling was achieved by the absence of glucagon and GLP-1 receptors (13; 14; 23), pharmacological action on the GLP-1R, or addition of exogenous GLP-1 or glucagon. Pharmacological inhibition and genetic deletion of receptors do not perturb the same signaling system. For example, ex-9 inhibits GLP-1Rs throughout the islet, including those expressed by δ-cells, whereas the conditional knockout selectively removes GLP-1R from β-cells. GLP-1R activation in δ-cells can promote somatostatin secretion, which in turn suppresses α-and β-cell activity (24–26). Thus, acute pan-islet GLP-1R inhibition may relieve δ-cell-mediated suppression of α-and β-cells (enhancing β-cell function) while simultaneously removing direct GLP-1R signaling from β-cells (inhibiting β-cell function). Therefore, it was important to perform these manipulations in the same study to compare the direct and indirect effects.

For comparison with the widely used isolated islet platform, we also utilized the pancreatic slice model, as the enzymatic digestion during islet isolation may affect the number of α-cells on the islet periphery. The pancreatic slice platform offers unperturbed α-cell numbers on the islet periphery, equal access of glucose to all cells in the imaging plane, and more physiologically relevant assessment of larger β-cell networks.

## Research Design and Methods

### Ethics Statement

Islet isolation from β-cell–specific GCaMP6s mice was approved by the University of Colorado IACUC (no:00024). Pancreatic slice experiments from C57BL/6J mice were approved by the Administration for Food Safety, Veterinary Sector and Plant Protection of the Republic of Slovenia (permit numbers U34401-35/2018-2). Isolated islet experiments from C57BL/6J mice were approved by the University of California IACUC (no:S23034). Islet isolation and pancreatic slice preparation were performed as previously described (6; 27).

To study the effect of the GLP-1R on islet function, we used tamoxifen-inducible β-cell specific humanized GLP-1R wild-type (WT) and knockout (KO) mice (28). KO was induced by oral tamoxifen (200mg/kg/day) for 3-4 days starting at 8 weeks of age. These experiments were approved by the University of California IACUC (no:23702).

### Glucose stimulation protocol and [Ca^2+^] imaging

Isolated islets and slices from non-GCaMP6s mice were stained with calbryte (AAT-Bioquest, 20653). One hour before imaging, samples were incubated in imaging solution containing BMHH and BSA (isolated islets) or HBS (for slices) and 2mM glucose. GCaMP6s islets and slices were imaged at 37°C with a 10min baseline at 2mM glucose, followed by 20min stimulation at 11mM, 20 min at 2mM, and a second 20min stimulation at 11 mM glucose. Calbryte-stained isolated islets were imaged for 30min in the 11mM glucose stimulations. In the experimental groups, 100nM exendin-9 (Sigma, E7269) or 100nM exendin-4 (Sigma, E7144) was added during the 2^nd^ part of the protocol (beginning of the second low glucose incubation).

GCaMP6s isolated islets were imaged on Zeiss LSM780 (40×) or LSM800 (20×) at 1Hz, ∼15µm from the islet surface. Pancreatic slices were imaged on Leica SP5 confocal systems (20x) at 2Hz, ∼15µm from the islet surface. Calbryte stained isolated islets were imaged using a Leica Stellaris 8 Falcon (20x) at 2 Hz, ∼13µm from the islet surface. A 488nm laser used to excite GCaMP6s/calbryte.

After live imaging, islets were fixed, permeabilized, blocked and stained with anti-insulin and anti-glucagon.

### Glucose uptake experiments

Isolated islets and slices were incubated for 1h in 2mM glucose imaging solution without 2-NBDG (ThermoFisher, N13195). Following a 1–2 min baseline, 2-NBDG was increased to 40µM (imaged using 488nm excitation), and in slices was washed out after 20min.

### Analysis of [Ca^2+^] dynamics

Cells were manually segmented from their [Ca^2+^] time series using ImageJ (FIJI) (29). Custom MATLAB routines were used to perform single-cell analysis of [Ca²□] dynamics. See supplementary information for more details.

*<u>1^st^ phase analysis</u>*: The peak height was defined as the most prominent local maximum during the 1^st^ phase response. Response time was defined as the time at which the [Ca²□] intensity reached 50% of the peak after the glucose increase.

*<u>Frequency analysis:</u>* After preprocessing, peaks were identified where the signal exceeded mean(residual) + 2×σ(residual) (30). Only cells for which var (signal) > var(noise) were used for analysis.

*<u>Network analysis:</u>* After pre-processing, Pearson correlation coefficients representing the temporal similarity between [Ca^2+^] traces were used to generate functional networks as previously described (21; 31–33).

### α-cell neighborhood

After background subtraction and median filtering, we aligned the IF images with the [Ca^2+^] time series. Each β-cell was represented using a sphere of diameter 25 μm, and the local α-cell neighborhood was calculated by quantifying the percent of glucagon-positive volume contained within the corresponding sphere, normalized to its volume.

### Statistical analysis

All statistical analysis was performed using GraphPad Prism (GraphPad, Boston, United States of America). Data are presented as box plots (median ± IQR) with min/max whiskers, and the mean indicated by a ‘+’. Differences were considered statistically significant for p < 0.05.

### Data availability statement

The datasets generated during and/or analyzed during the current study are available from the corresponding author upon reasonable request.

### Resource availability statement

The MATLAB codes generated during and/or analyzed during the current study is available from the corresponding author upon reasonable request.

## Results

In previous studies, the manipulation of α-β signaling was achieved by the absence of glucagon and GLP-1 receptors (13; 14; 23), or addition of exogenous glucagon/GLP-1. In contrast, we stimulated the islet with glucose consecutively while also pharmacologically manipulating the GLP-1 receptors with ex4/ex-9 during the second stimulation. This approach, while technically more complex, allowed us to compensate for inter-islet heterogeneity, and directly compare changes in the functional subpopulations and their α-cell proximity.

### GLP-1R potentiates 1^st^-phase response to glucose

We compared 1^st^ glucose stimulation with the 2^nd^ stimulation in the presence or absence of ex-9/ex-4 (**Fig. 1A,B** and **S1A,C**). In both platforms, in control and ex-4 groups, the response time, T_resp_, became shorter in the 2^nd^ stimulation (**Fig. 1C,G**) compared to 1^st^ stimulation. After compensating for the heterogeneity by subtracting T_resp_ in the 2^nd^ stimulation from the T_resp_ in 1^st^ stimulation, we found that in both platforms, GLP-1R inhibition significantly prolonged the T_resp_ compared to control and agonist groups (**Fig. 1D,H**). Effects of the GLP-1R agonist, ex-4 were inconsistent between the platforms.

**Figure 1.**
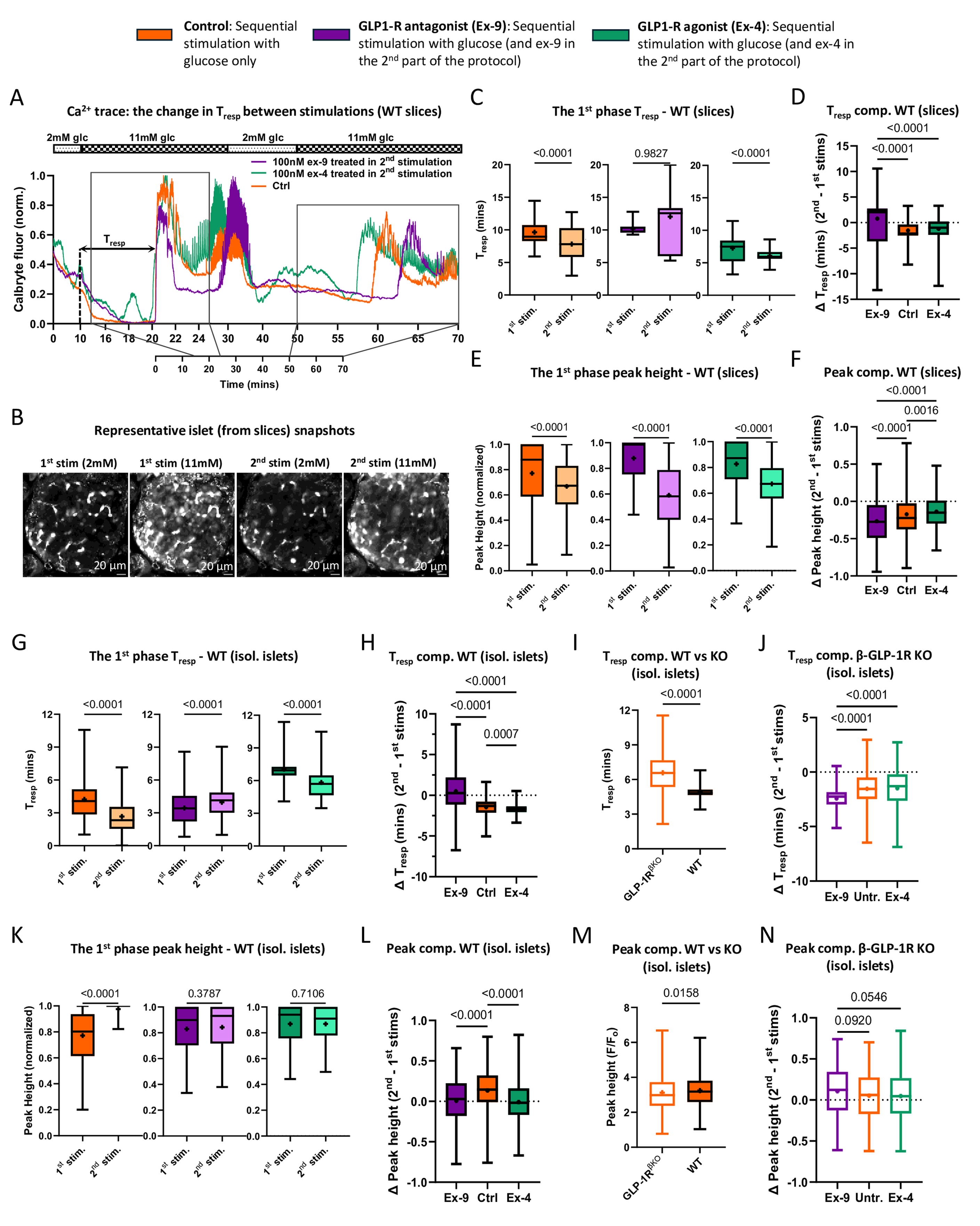
Effects of GLP-1R signaling on the β-cell 1^st^ phase [Ca²□] response to glucose. **(A)** Experimental protocol for the different treatments with representative islet-average [Ca²□] time courses highlighting the 1^st^ phase response in slices. Traces were aligned to the onset of the 1^st^ glucose-induced 1^st^ phase response to facilitate visual comparison of the 2^nd^ stimulation. **(B)** Representative islet snapshots at selected time points from slices. Time of response to glucose of each β-cell in 1^st^ vs 2^nd^ stimulations in slices **(C)** and isolated islets **(G)** (slices: n = 9 control, n = 6 ex-9, and n = 5 ex-4 treated islets; isolated islets: n = 18 control, n = 18 ex-9, and n = 7 ex-4 treated islets). Comparison of change in response time quantified as differences in response time (2^nd^ – 1^st^ stimulations) in slices **(D)** and isolated islets **(H).** Negative values indicate faster responses in the 2^nd^ stimulation, and positive values indicate slower responses. 1^st^ phase peak height of each β-cell in 1^st^ vs 2^nd^ stimulations in slices **(E)**, and isolated islets **(K)** (slices: n = 6 control, n = 5 ex-9, and n = 5 ex-4 treated islets; isolated islets: n = 8 control, n = 7 ex-9, and n = 7 ex-4 treated islets). Comparison of difference in the peak height (2^nd^ – 1^st^ stimulations) in slices **(F)** and isolated islets **(L)**. **(I)** Time of response to glucose of each β-cell in WT vs KO islets (n = 6 WT and n = 26 KO islets). **(J)** Comparison of change in response time (2^nd^ – 1^st^ stimulations) in KO islets (n = 9 untreated, n = 7 ex-9 treated and n = 10 ex-4 treated islets). **(M)** 1^st^ phase peak height of each β-cell in WT vs KO islets (n = 6 WT and n = 28 KO islets). **(N)** Comparison of difference in the peak height (2^nd^ – 1^st^ stimulations) in KO islets (n = 9 untreated, n = 9 ex-9 treated and n = 10 ex-4 treated islets). Outliers were removed using the ROUT method (Q = 1%). Wilcoxon matched-pairs signed-rank test was used for statistical analysis in panels **(C)**, **(E) (G)** and **(K).** Kruskal-Wallis test with Dunn’s multiple comparison tests was used for statistical analysis in panels **(D)**, **(F)**, **(H)**, **(J)**, **(L)** and **(N)**. Mann Whitney test was used for statistical analysis in panels **(I)** and **(M)**.

To complement the pharmacological studies, we utilized isolated islets from β-cell-specific humanized GLP-1R KO mice. In the absence of GLP-1Rs on β-cells, islets exhibited a longer response time (**Fig**. **1I**), in agreement with the effects of ex-9 treatment of the WT islets (**Fig. 1H**). In the KO islets we also observed shorter response time with ex-9 treatment - an off-target effect (**Fig. 1J**), potentially reflecting altered δ-cell signaling, signaling through the glucagon receptor, or other pathways. These possibilities require direct experimental testing.

We next compared the 1^st^-phase peak amplitude in the 1^st^ vs 2^nd^ glucose stimulation (**Fig. 1E,K**). Again, after compensating for the inter-sample heterogeneity, in both models, we found a greater loss of the peak upon GLP-1R inhibition, compared to control group (**Fig. 1F,L**). The KO islets also had lower peak amplitude compared to the WT (**Fig. 1M**). We found no off-target effects of ex-9 or ex-4 on the peak amplitude in the KO islets (**Fig. 1N**). Overall, these findings suggest the role of GLP-1R signaling in enhancing and accelerating the 1^st^ phase [Ca^2+^] response.

### GLP-1R signaling promotes the spatiotemporal consistency of 1^st^ responders and their α-cell association in pancreatic slices

After recording [Ca^2+^] dynamics in the live islets **(Fig. 2A)**, we then fixed and stained these islets for insulin and glucagon (**Fig. 2B** and **S1D**).

**Figure 2.**
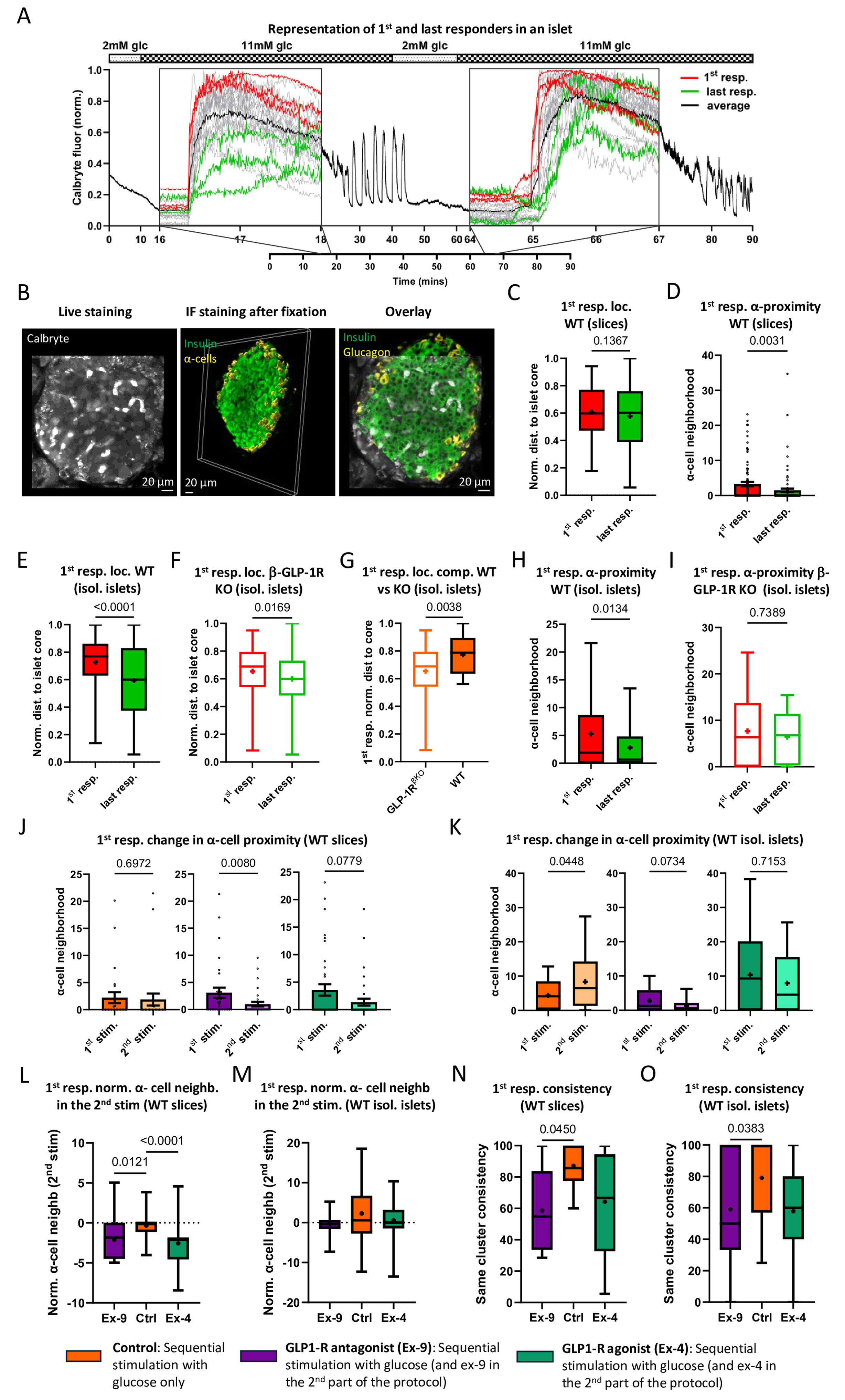
Effects of GLP-1R signaling on 1^st^ responder identity and spatial organization in the islet. **(A)** Representative [Ca²□] time course from an isolated islet, with the first (red) and last (green) responders highlighted. **(B)** Representative calbryte-stained live slice used for [Ca²□] time-series; 3D rendering of the IF z-stack after fixation; live and fixed image overlays. Comparison of the location of 1^st^ and last responders in the islet relative to the center of the islet in slices **(C)** and isolated islets **(E)** (slices: n = 15 islets; isolated islets: n = 42 islets). Value 1 corresponds to the islet surface and 0 to the islet core (center of the optical 2D plane). Comparison of α-cell neighborhood in the immediate proximity of 1^st^ and last responders in slices **(D)** and isolated islets **(H)** (slices: n = 12 islets; isolated islets: n = 19 islets). **(F)** Comparison of the location of 1^st^ and last responders in the KO islets relative to the center of the islet (n = 28 islets). **(G)** Comparison of location of 1^st^ responders in the islet (relative to the center of the islet) between WT and KO islets (n = 6 WT and n = 28 KO islets). **(I)** Comparison of α-cell neighborhood in the immediate proximity of 1^st^ and last responders in KO islets (n = 8 islets). α-cell neighborhood in the immediate proximity of 1^st^ responders independently identified in the 1^st^ vs 2^nd^ stimulations in slices **(J)** and isolated islets **(K)** (slices: n = 5 control, n = 4 ex-9, and n = 4 ex-4 treated islets; isolated islets: n = 9 control, n = 6 ex-9, and n = 5 ex-4 treated islets). Comparison of change in 1^st^ responder α-cell neighborhood quantified as the normalized 2^nd^ stimulation α-cell neighborhood (normalized by subtracting each islet’s mean 1^st^ responder α-cell neighborhood during the 1^st^ stimulation) in slices **(L)** and isolated islets **(M).** Values lower than 0 indicate a decrease in the 2^nd^ stimulation. Comparison of consistency of 1^st^ responders: percent of the cells identified as a 1^st^ responder in the 2^nd^ stimulation that were also a 1^st^ responder, 1^st^ nearest neighbor, or 2^nd^ nearest neighbor (same cell cluster) in the 1^st^ stimulation in slices **(N)** and isolated islets **(O)** (slices: n = 9 control, n = 6 ex-9, and n = 5 ex-4 treated islets; isolated islets: n = 24 control, n = 15 ex-9, and n = 7 ex-4 treated islets). Outliers were removed using the ROUT method (Q = 1%). Mann-Whitney test was used for statistical analysis in panels **(C)** - **(K)**, **(N)** and **(O)**. Kruskal-Wallis test with Dunn’s multiple comparison tests was used for statistical analysis in panels **(L)** and **(M)**.

The 40μm-thick volumetric immunofluorescence images were collected and registered to the corresponding live images within the same focal plane. 1^st^ responders were identified as a functional subpopulation of β-cells that drive the islet’s 1^st^ phase [Ca^2+^] response (6). The 1^st^ responders were located significantly closer to the islet periphery than the last responders in isolated islets, but not in slices (**Fig. 2C,E**). The β-cell-GLP-1R-KO caused a decrease in significance of the difference between 1^st^ vs last responders (**Fig. 2F**). Moreover, 1^st^ responders in the KO model were further from the islet periphery than in the WT islet (**Fig. 2G**). In both platforms, the 1^st^ responders had significantly higher number of α-cell neighbors compared to the last responders (**Fig. 2D**,**H**), and this 1^st^-responder-α-cell association was disrupted by the KO (**Fig. 2I**). In slices (but not in the isolated islets) we observed a significant decrease in the α-cell proximity of 1^st^ responders upon GLP-1R inhibition **(Fig. 2J-M)**. Additionally, in both platforms, GLP-1R inhibition resulted in a significant decrease of the 1^st^ responder consistency **(Fig. 2N,O)**. These results suggest that GLP-1R signaling plays a role in maintaining the spatiotemporal consistency of 1^st^ responders, and that its action depends on α-cell proximity.

### Glucose penetration into pancreatic slices differs from that of isolated islets

To test whether some of the differences observed between pancreatic slices and isolated islets resulted from differences in glucose delivery to cells in the islet core, we monitored the penetration of the fluorescent glucose analog 2-(N-(7-Nitrobenz-2-oxa-1,3-diazol-4-yl)Amino)-2-Deoxyglucose over time in both platforms **(Fig. 3A,D)**. Representative images of 2-NBDG fluorescence at selected timepoints for the slices and islets are shown in **(Fig. 3B,E)**. Following imaging, we identified the different endocrine cell-types (**Fig. 3C,F**). For each β-cell, we plotted the time required for 2-NBDG fluorescence to reach 25% of the fluorescence intensity in the surrounding medium against the cell’s distance from the islet periphery **(Fig. 3G,H).** There was a significantly higher gradient of the 2-NBDG penetration into isolated islets (4.5 sec/μm) compared to slices (0.5 sec/μm), corresponding to apparent speed of the glucose uptake of 13.2 μm/min in isolated islets and 120 μm/min in slices **(Fig. 3I)**. This was likely due to almost immediate access of the glucose to all cells in the dissected islets within the slice, as opposed to non-dissected isolated islets. These findings offer a plausible explanation of the larger difference in 1^st^ vs last responder’s location in the isolated islets, compared to the slices.

**Figure 3.**
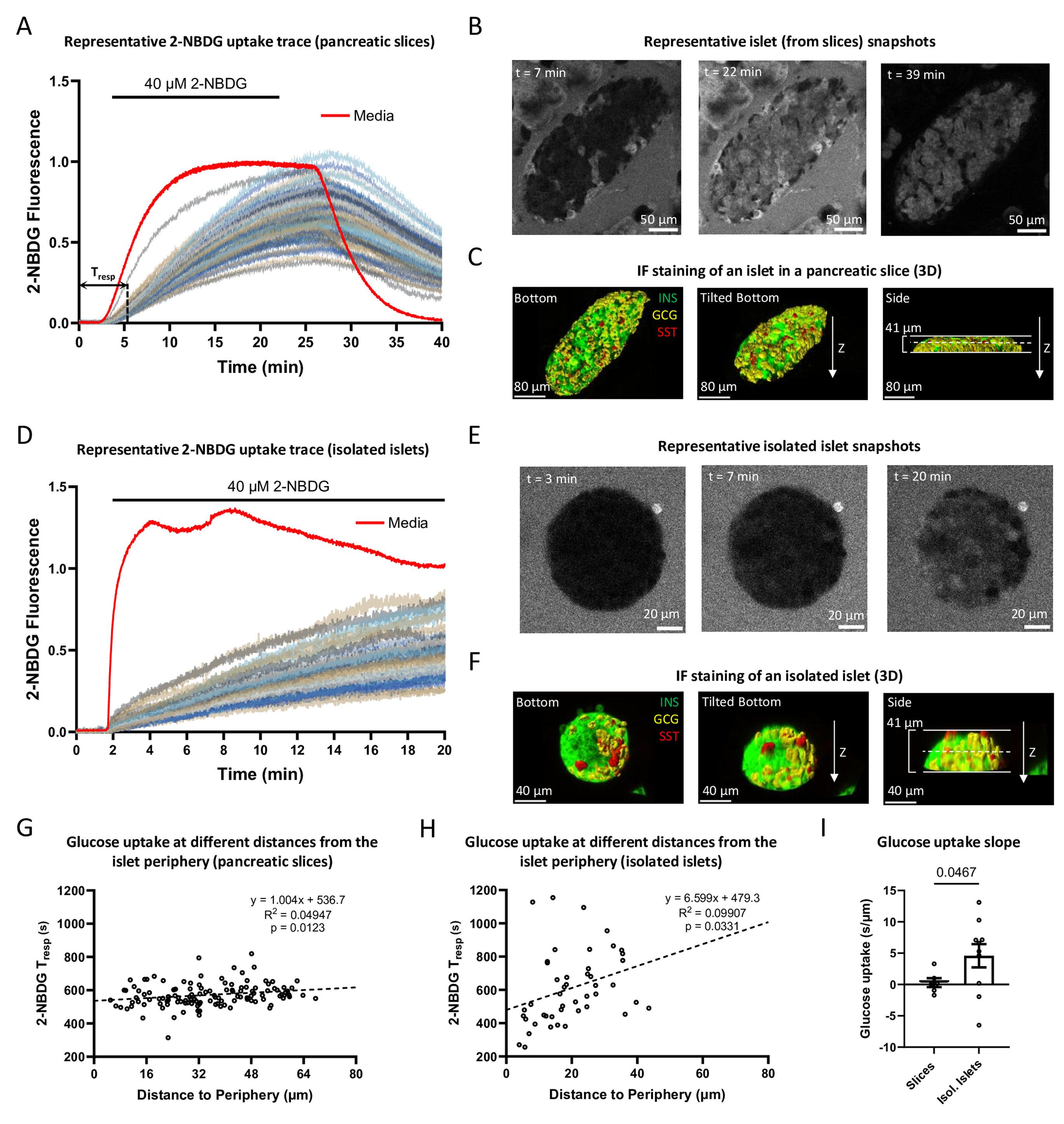
Comparison of glucose uptake dynamics between isolated islets and pancreatic slices. Experimental protocol and representative 2-NBDG fluorescence time courses showing glucose uptake dynamics in slices **(A)** and isolated islets **(D)**. Representative islet snapshots at selected time points of the experiments from slices **(B)** and isolated islets **(E)**. 3D renderings of IF z-stacks from representative islets following fixation after live imaging of slices **(C)** and isolated islets **(F)**. Relationship between 2-NBDG response time and distance from the islet periphery in a representative pancreatic slice **(G)** and a representative isolated islet **(H)**. Linear regression slopes were used to quantify glucose uptake dynamics in the islet. **(I)** Comparison of glucose uptake slopes between isolated islets and islets in intact slices (n = 6 slices and n = 10 isolated islets). Simple linear regression was used for statistical analysis in panels **(G)** and **(H)**. Mann Whitney test was used for statistical analysis in panel **(I)**.

### GLP-1R communication increases the speed of [Ca^2+^] signal propagation

Since β-cell responses often occur within localized clusters or domains (34; 35), we created an automated, bias-free ‘1^st^ responder domain’ recognition algorithm as shown in **Fig. 4A-C** (see **Fig. S2)**. Each domain consists of a local 1^st^ responder initiating the [Ca^2+^] signal during the 1^st^ phase of the islet’s response to glucose, propagating to all the other β-cells in the domain. Within each domain we plot the distance of each β-cell to the local 1^st^ responder vs β-cell’s response time. The slope of this graph indicated the apparent speed of the [Ca^2+^] signal for that domain (**Fig. 4D**). In both platforms the speed of signal propagation in the 2^nd^ stimulation significantly dropped upon ex-9 addition, compared to the 1^st^ stimulation (**Fig. 4E,H**). We observed a mild decrease in speed for the GLP-1R inhibition group compared to control **(Fig. 4F,I)**, consistent with the KO islets, where the speed was significantly lower than in the WT (**Fig. 4G**). No significant changes in speed upon ex-9/ex-4 treatment of the KO islets were observed, however a trend towards diminished speed under ex-9 was observed, suggesting a possible indirect effect (**Fig. 4J**).

**Figure 4.**
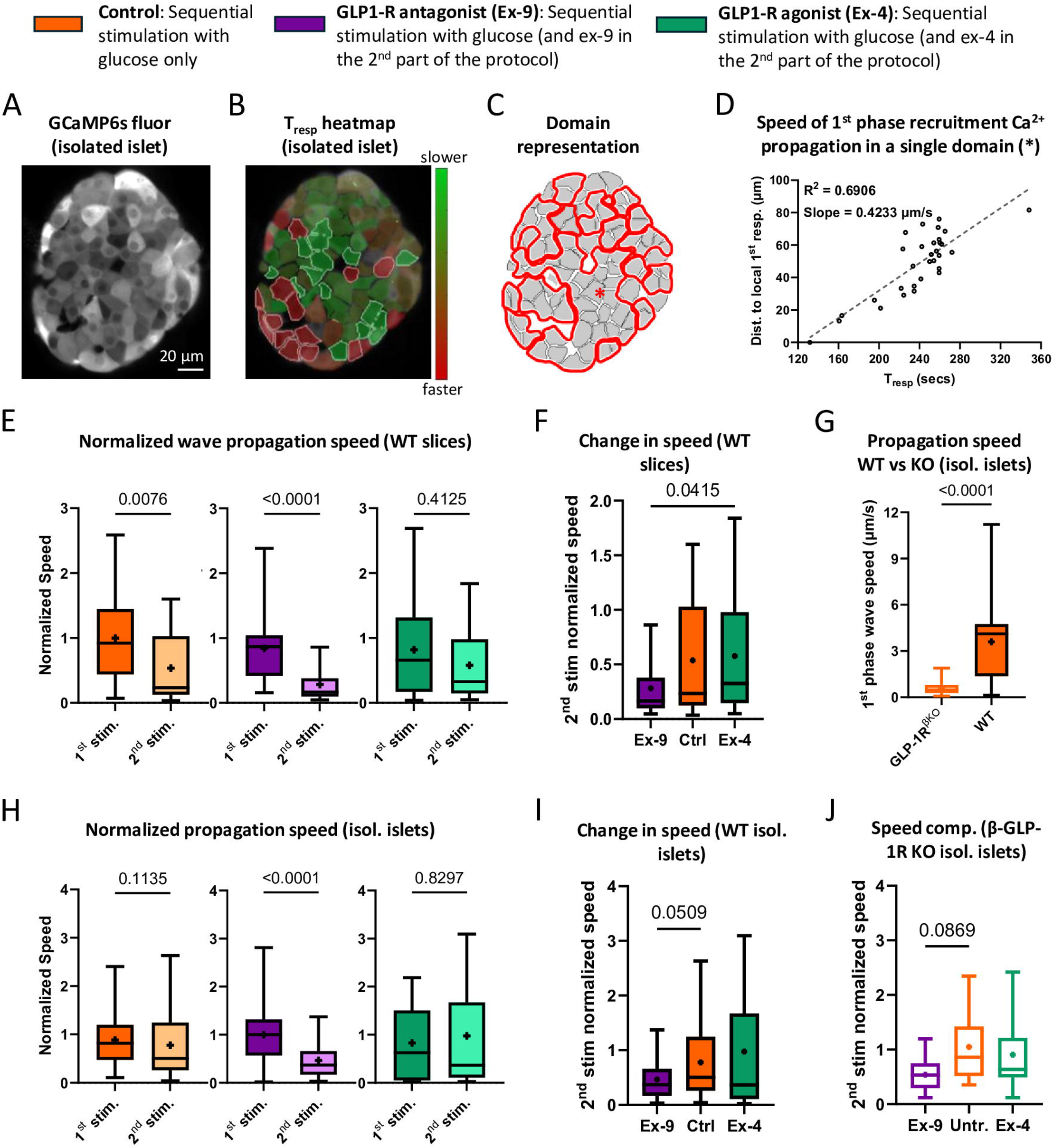
Effects of GLP-1R signaling on speed of excitation propagation in β-cells during the 1^st^ phase [Ca^2+^] response to glucose. **(A)** Representative [Ca²□] image of a GCaMP6s expressing isolated islet. **(B)** Corresponding β-cells pseudo-colored based on their response time to glucose. **(C)** Corresponding response time domains, each defined by a local 1^st^ responder. Domain boundaries were identified based on the linear dependence between the response time vs distance to the local 1^st^ responder, as illustrated for a representative domain (indicated by a *) in **(D).** [Ca^2+^] signal wave propagation speed of each domain (normalized to each islet’s mean domain speed during the 1^st^ stimulation) in 1^st^ vs 2^nd^ stimulations in slices **(E)** and isolated islets **(H)** (slices: n = 7 control, n = 5 ex-9, and n = 5 ex-4 treated islets; isolated islets: n = 18 control, n = 18 ex-9, and n = 6 ex-4 treated islets). Comparison of 2^nd^ stimulation normalized speed (representation of change in speed) in slices **(F)** and isolated islets **(I)**. **(G)** [Ca^2+^] signal wave propagation speed of each domain in WT vs KO islets (n = 6 WT and n = 28 KO islets). Values lower than 1 indicate a decrease in speed in the 2^nd^ stimulation. **(J)** Comparison of the 2^nd^ stimulation normalized speed in KO islets (n = 9 untreated, n = 8 ex-9 treated and n = 10 ex-4 treated islets). Outliers were removed using the ROUT method (Q = 1%). Simple linear regression was used for statistical analysis in panel **(D)**. Mann Whitney test was used for statistical analysis in panels **(E)**, **(G)** and **(H)**. Kruskal-Wallis test with Dunn’s multiple comparison tests was used for statistical analysis in panels **(F)**, **(I)** and **(J)**.

### GLP-1R signaling regulates calcium oscillation frequency and active time, especially in the α-neighboring β-cells

We applied ex-9 and ex-4 as before (**Fig. 5,6 A**). The images of fluorescent-indicator-stained islets, pseudo color maps of the frequency change, and IF in the same focal plane are shown in **Fig. 5C,F,I** and **6D,G,J**. In control and ex-4 groups, the frequency increased relative to the 1^st^ stimulation, in both platforms. For the ex-9 group, this effect was significantly diminished or reversed (**Fig. 5D,G,J** and **6E,H,K**). After compensating for islet heterogeneity, we observed highly significant differences between all groups in both platforms. Namely, the frequency increases were much higher upon GLP-1R agonism, and lower upon GLP-1R antagonism, relative to the control group **(Fig. 5,6 B)**. In the KO islets there was no difference between the groups (**Fig. 6C**).

**Figure 5.**
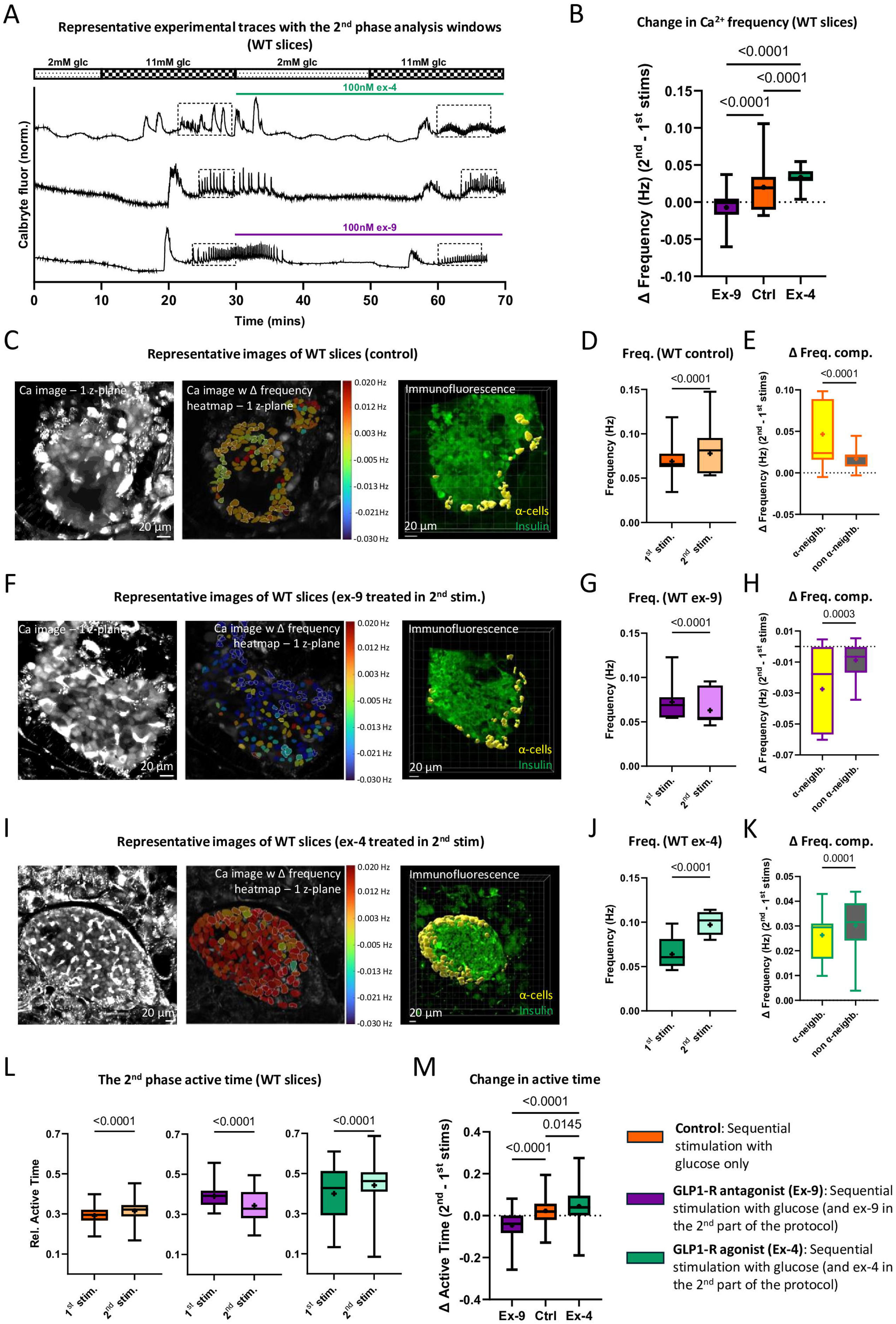
Effects of GLP-1R signaling on the β-cell 2^nd^ phase [Ca²□] frequency in pancreatic slices. **(A)** Representative islet-average [Ca²□] traces for the different treatments and the analysis windows (boxed) used for frequency and functional network analysis during the 1^st^ and 2^nd^ stimulations. **(B)** Comparison of the change in oscillation frequency (2^nd^ – 1^st^ stimulations) between control, ex-9-, and ex-4-treated islets, quantified from **(D)**, **(G)**, and **(J)**. Time-projection of [Ca^2^] time series; heatmap representing change in oscillation frequency between two stimulations; z-projection of the IF image with α-cells segmented in Imaris in a representative islet for a control **(C)**, ex-9-treated **(F)**, and ex-4-treated **(I)** islets. Oscillation frequency of each β-cell in 1^st^ vs 2^nd^ stimulations in control **(D)**, ex-9-treated **(G)**, and ex-4-treated **(J)** islets (n = 9 control, n = 5 ex-9, and n = 5 ex-4 treated islets). Comparison of change in oscillation frequencies (2^nd^ – 1^st^ stimulations) between α-neighboring and non α-neighboring β-cells in control **(E)**, ex-9-treated **(H)**, and ex-4-treated **(K)** islets (n = 6 control, n = 4 ex-9, and n = 4 ex-4 treated islets). **(L)** Relative active time of each β-cell in 1^st^ vs 2^nd^ stimulations in control, ex-9-treated, and ex-4-treated islets (n = 5 control, n = 5 ex-9, and n = 5 ex-4 treated islets). **(M)** Comparison of change in active time (2^nd^ – 1^st^ stimulations) between control, ex-9-, and ex-4-treated islets. Outliers were removed using the ROUT method (Q = 1%). Kruskal-Wallis test with Dunn’s multiple comparison tests was used for statistical analysis in **(B)** and **(M).** Wilcoxon matched-pairs signed-rank test was used for statistical analysis in panels **(D)**, **(G)**, **(J),** and **(L)**. Mann Whitney test was used for statistical analysis in panels **(E)**, **(H)**, and **(K)**.

**Figure 6.**
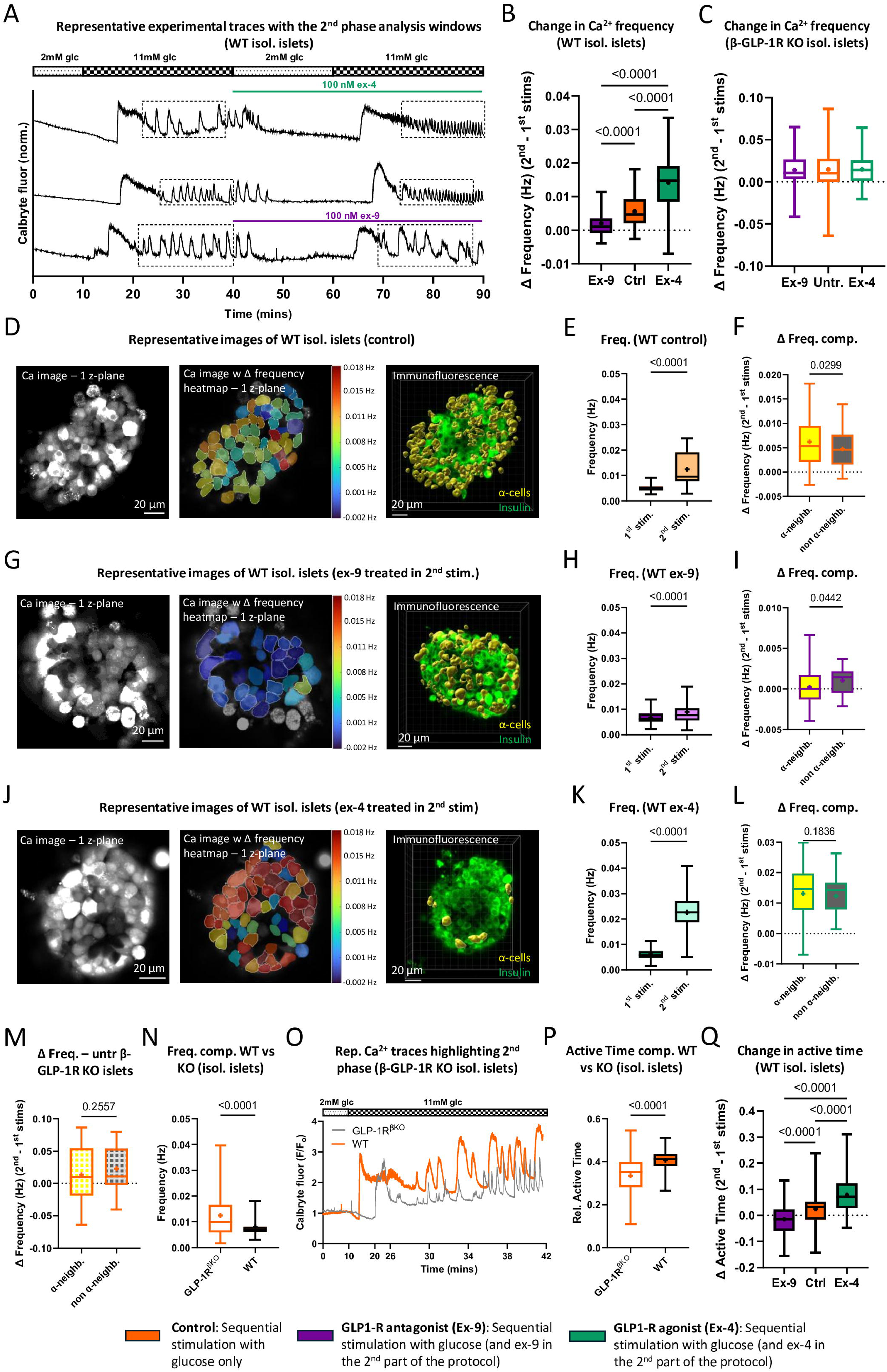
Effects of GLP-1R signaling on the β-cell 2^nd^ phase [Ca²□] frequency in isolated islets. **(A)** Representative islet-average [Ca²□] traces for the different treatments and the analysis windows (boxed) used for frequency and functional network analysis during the 1^st^ and 2^nd^ stimulations. **(B)** Comparison of the change in oscillation frequency (2^nd^ – 1^st^ stimulations) between control, ex-9-, and ex-4-treated islets, quantified from **(E)**, **(H)**, and **(K)**. **(C)** Comparison of change in oscillation frequency (2^nd^ – 1^st^ stimulations) in KO islets (n = 9 untreated, n = 9 ex-9 treated and n = 10 ex-4 treated islets). Time-projection of [Ca^2^] time series; heatmap representing change in oscillation frequency between two stimulations; z-projection of the IF image with α-cells segmented in Imaris in a representative islet for a control **(D)**, ex-9-treated **(G)**, and ex-4-treated **(J)** islets. Oscillation frequency of each β-cell in 1^st^ vs 2^nd^ stimulations in control **(E)**, ex-9-treated **(H)**, and ex-4-treated **(K)** islets (n = 8 control, n = 6 ex-9, and n = 7 ex-4 treated islets). Comparison of change in oscillation frequencies (2^nd^ – 1^st^ stimulations) between α-neighboring and non α-neighboring β-cells in control **(F)**, ex-9-treated **(I)**, and ex-4-treated **(L)** islets (n = 7 control, n = 4 ex-9, and n = 5 ex-4 treated islets). **(M)** Comparison of change in oscillation frequencies (2^nd^ – 1^st^ stimulations) between α-neighboring and non α-neighboring β-cells in untreated KO islets (n = 4 islets). **(N)** 2^nd^ phase oscillation frequency of each β-cell in WT vs KO islets (n = 5 WT and n = 28 KO islets). **(O)** Representative islet average [Ca]^2+^ traces from WT and KO islets highlighting the 2^nd^ phase. **(P)**. 2^nd^ phase relative active time of each β-cell in WT vs KO islets (n = 6 WT and n = 9 KO islets). **(Q)** Comparison of change in active time (2^nd^ – 1^st^ stimulations) between control, ex-9-, and ex-4-treated islets (n = 5 control, n = 5 ex-9, and n = 5 ex-4 treated islets). Outliers were removed using the ROUT method (Q = 1%). Kruskal-Wallis test with Dunn’s multiple comparison tests was used for statistical analysis in **(B)**, **(C)** and **(Q).** Wilcoxon matched-pairs signed-rank test was used for statistical analysis in panels **(E)**, **(H)** and **(K)**. Mann Whitney test was used for statistical analysis in panels **(F)**, **(I)**, **(L)**, **(M)**, **(N)** and **(P)**.

In most islets with a higher α-cell count, we observed a higher oscillatory frequency (**Fig. S3A**), in agreement with previously published findings (36). After aligning the [Ca^2+^] time series recordings with their corresponding volumetric IF images we observed a significantly greater increase in frequency in α-neighboring β-cells compared to non-α-neighboring cells in control experiments in both platforms (**Fig. 5E** and **6F**). Upon GLP-1R inhibition, this effect was diminished or reversed – more so in the α-neighboring β-cells in both platforms (**Fig. 5H** and **6I**). The difference between α-neighboring and non-α-neighboring β-cells was erased by the KO (**Fig. 6M**). These findings suggest that α-cells exert localized influence on β-cell frequency of [Ca²□] oscillations during the 2^nd^ phase of the response via GLP-1R mediated pathway.

### Impaired active time, but not frequency was found in the β-cell-specific GLP-1R KO

Surprisingly, we found that islets from the KO mice had higher frequency of Ca^2+^ oscillations compared to the islets in the WT (**Fig. 6N**). However, after a closer examination of the traces, we found that the higher frequency did not translate into the gain-of-function effect (**Fig. 6O**). On the contrary, the KO islets exhibited a lower active time (**Fig. 6P**). We then accessed the active time in the control, ex-9, and ex-4 groups in the WT islets and found a gain-of-function under ex-4 and a loss-of-function under ex-9, compared to control, in both platforms (**Fig. 5L,M, S3B** and **6Q**).

### GLP-1R signaling enhances the functional connectivity

We generated functional β-cell networks for the 1^st^ and 2^nd^ stimulations **(Fig. 7A** and **S3D)**. After compensation for islet heterogeneity, in both platforms ex-4 resulted in a significant increase in connectivity relative to control group **(Fig. 7B,C,E,F)**. Surprisingly, the same effect was observed for ex-9 group, unlike the loss of connectivity in the KO (**Fig. 7D**). Pharmacological manipulation of the KO islets yielded no difference between the groups (**Fig. 7G**), indicating no off-target effects.

**Figure 7.**
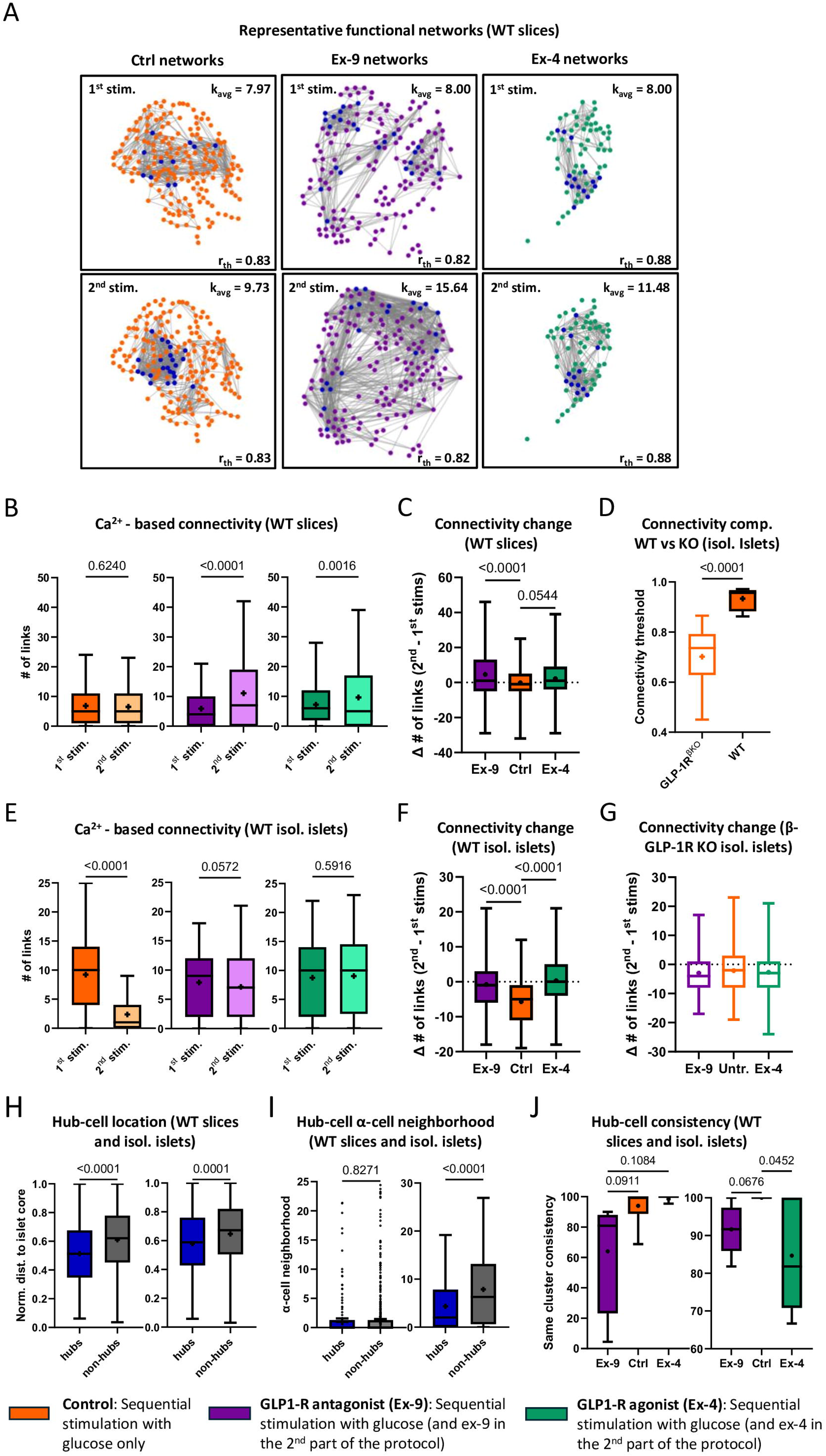
Effects of GLP-1R signaling on the β-cell 2^nd^ phase [Ca²□] functional connectivity. **(A)** Functional β-cell networks in the 1^st^ and 2^nd^ stimulations for representative control, ex-9 treated and ex-4 treated slices. Functional connectivity of each β-cell in 1^st^ vs 2^nd^ stimulations in slices **(B)** and isolated islets **(E)** (slices: n = 7 control, n = 4 ex-9, and n = 4 ex-4 treated islets; isolated islets: n = 8 control, n = 5 ex-9, and n = 5 ex-4 treated islets). Comparison of change in functional connectivity (2^nd^ – 1^st^ stimulations) in slices **(C)** and isolated islets **(F)**. **(D)** 2^nd^ phase connectivity threshold in WT vs KO islets required to achieve an average of 8 links per cell (n = 6 WT and n = 18 KO islets). **(G)** Comparison of change in connectivity, quantified as the number of links (2^nd^ – 1^st^ stimulations) in KO islets (n = 6 untreated, n = 6 ex-9 treated and n = 6 ex-4 treated islets). **(H)** Comparison of the location of hub-cells and non-hub cells in the islet relative to the center of the islet (slices: n = 15 islets; isolated islets: n = 18 islets). Value 1 corresponds to the islet surface and 0 to the islet core (center of the optical 2D plane). **(I)** Comparison of α-cell neighborhood in the immediate proximity of hub cells and non-hub cells (slices: n = 10 islets; isolated islets: n = 18 islets). **(J)** Comparison of consistency of hub cells: percent of the cells identified as a hub in the 2^nd^ stimulation that were also a hub, 1^st^ nearest neighbor, or 2^nd^ nearest neighbor (same cell cluster) in the 1^st^ stimulation (slices: n = 6 control, n = 4 ex-9, and n = 3 ex-4 treated islets; isolated islets: n = 8 control, n = 5 ex-9, and n = 5 ex-4 treated islets). Outliers were removed using the ROUT method (Q = 1%). Wilcoxon matched-pairs signed-rank test was used for statistical analysis in panels **(B)** and **(E)**. Kruskal-Wallis test with Dunn’s multiple comparison tests was used for statistical analysis in panels **(C)**, **(F)**, **(G)** and **(J)**. Mann Whitney test was used for statistical analysis in panels **(D)**, **(H)** and **(I)**.

### Hub-cells consistency is maintained via islet-wide GLP-1R signaling

Hub-cells were identified as functional β-cell subpopulation that coordinate β-cell [Ca^2+^] oscillations in the islet (7), (37). In both platforms they were located significantly closer to the islet core compared to non-hubs (**Fig. 7H**). In isolated islets, hub cells were significantly further away from the α-cells (**Fig. 7I**). Finally, addition of ex-9 disrupted spatial consistency of hub cells in both platforms (**Fig. 7J**). Together these results suggest that consistency of hub cells is not dependent on the distance to α-cells, but more on the islet-wide GLP-1R-mediated effects.

## Discussion

We demonstrate that the islet-endogenous GLP-1R signaling is an important regulator of both the first and second phases of the β-cell response to glucose. It does not only amplify overall β-cell activity but contributes to the temporal and spatial organization of β-cell networks. The phenotype produced by β-cell-specific GLP-1R KO was broadly consistent with GLP-1R inhibition in wild-type islets. This suggests that constitutive or locally generated islet GLP-1R signaling may be sufficient to influence β-cell function under glucose stimulation.

### GLP-1R signaling amplifies both phases of the β-cell glucose response

During the 1^st^ phase, inhibition of GLP-1R signaling prolonged the β-cell’s time of response to glucose, reduced the peak amplitude, and reduced the apparent speed of Ca² signal propagation through β-cell domains, consistent with the similar effects produced by β-cell-specific GLP-1R KO. These observations indicate that GLP-1R signaling increases the readiness of β-cells to respond to glucose and facilitate recruitment of neighboring cells during the initial response to glucose, rather than acting exclusively at the level of insulin granule exocytosis.

The effects on the 2^nd^ phase were complementary. Under control conditions, Ca² oscillation frequency increased during the second glucose stimulation, resembling the well-described time-dependent potentiation or “metabolic memory” of insulin secretion, previously-reported under consecutive glucose stimulation (38), attributed to CaMKII, protein kinase C, and other mechanisms (39–41). Therefore, the absolute difference between the first and second stimulation cannot be attributed solely to GLP-1R signaling. Our within-islet design nevertheless allowed us to distinguish the additional effect of GLP-1R manipulation by comparing the change between stimulations across experimental groups. GLP-1R inhibition reduced the oscillation frequency and active time, whereas GLP-1R agonism enhanced it, compared to the control group. The β-cell-specific GLP-1R knockout further supported a role for GLP-1R signaling in maintaining sustained β-cell activity, although the knockout phenotype was more apparent in active time than in oscillation frequency.

Together, these findings are consistent with the established ability of GLP-1R signaling to increase cAMP and activate PKA and Epac2A pathways that potentiate β-cell stimulus-secretion coupling (42; 43). Increased cAMP signaling can enhance the efficiency of the secretory machinery and modulate ion-channel activity, providing plausible mechanisms through which GLP-1R activation could alter both Ca² dynamics and insulin secretion (44–47). Direct measurements of cAMP together with Ca² dynamics will be required to determine whether the effects observed here are mediated by changes in cAMP signaling.

### α-cells proximity links GLP-1R signaling to spatially heterogeneous β-cell function

First-responder β-cells, which initiate the first-phase Ca² response, had a greater local abundance of α-cells than last responders. Moreover, GLP-1R inhibition disrupted the spatial and temporal consistency of first responders, suggesting that functional β-cell heterogeneity is not determined exclusively by cell-intrinsic properties. Instead, the local endocrine environment may contribute to establishing or maintaining functional β-cell states. Glucagon and other proglucagon-derived peptides can activate β-cell cAMP signaling through GLP-1R and glucagon receptors (13; 14; 23; 48–50), providing a potential mechanism for local α-cell modulation of β-cell activity. Because α-cell-derived signals are expected to be spatially concentrated around their site of release, β-cells near α-cells may experience a different paracrine environment from β-cells located farther away.

β-cells neighboring α-cells showed a greater increase in oscillation frequency during repeated glucose stimulation, and this enhancement was diminished or reversed by GLP-1R inhibition. The KO resulted in loss of the functional difference between the α-neighboring and non-α-neighboring β-cells. These findings are consistent with previous observations that α-and β- cell activity is coordinated within intact islets (36) and provide a potential functional mechanism through which α-cells can contribute to β-cell heterogeneity. Future experiments combining cell-specific manipulation of glucagon/proglucagon signaling with simultaneous measurements of cAMP and Ca² will be important for resolving this mechanism.

### GLP-1R signaling contributes to the functional organization of β-cell networks

The effects of GLP-1R signaling extended beyond individual β-cell responses to the organization of multicellular Ca² activity. First responders initiate local Ca² activity, while hub cells coordinate activity across larger portions of the β-cell network (6–8). GLP-1R inhibition reduced the speed of signal propagation within first-responder domains and disrupted the spatial consistency of hub cells. β-cell-specific GLP-1R deletion also reduced functional connectivity relative to wild-type islets. Together, these findings suggest that GLP-1R signaling contributes not only to the magnitude of β-cell responses but also to how these responses are distributed through the islet network.

The distinction between first responders and hub cells is important. First responders are spatially associated with α-cells and are involved primarily in initiation of the first-phase response, whereas hub cells represent a network-level population associated with coordination of Ca² oscillations. The differential spatial relationship of these populations to α-cells suggests that GLP-1R signaling may influence β-cell network organization through both local and islet-wide mechanisms. Local α-cell signaling may preferentially influence initiation and oscillatory behavior in neighboring β-cells, whereas GLP-1R signaling throughout the β-cell population may help maintain the network properties required for coordinated activity.

An unexpected finding was that acute ex-9 treatment increased functional connectivity in WT islets, whereas genetic loss of β-cell GLP-1R reduced connectivity. This could be due to pan-islet ex-9 treatment resulting in competing effects on different endocrine cell populations. Importantly, however, the present experiments do not directly measure somatostatin or glucagon secretion. Therefore, the proposed δ-cell mechanism should be considered a testable hypothesis.

### Physiological relevance and limitations

Our findings demonstrate that GLP-1R signaling contributes to the organization of β-cell function under normal glucose stimulation, rather than merely serving as a pharmacologically activated pathway. The islet does not respond to glucose as a homogeneous population of β-cells. Instead, distinct cells initiate, propagate, and coordinate Ca² activity, while their functional behavior is influenced by the surrounding endocrine microenvironment. Endogenous α-β communication helps maintain both rapid recruitment of β-cells and sustained oscillatory activity.

The comparison between isolated islets and pancreatic slices is important in this context. Islet isolation can alter the native spatial relationship between endocrine cells, particularly at the islet periphery, whereas pancreatic slices preserve a more intact local α-cell environment. Consistent with this, several effects involving α-cell proximity were stronger or more apparent in slices. Our glucose-penetration experiments further demonstrated that glucose access is less efficient in the isolated islets, cautioning against directly extrapolating the magnitude of an effect observed in isolated islets to the intact pancreas.

Several limitations should be considered. First, Ca² imaging provides a sensitive readout of β-cell electrical activity but is not a direct measurement of insulin secretion. Although the measured parameters are closely related to stimulus-secretion coupling, future studies combining single-cell Ca² imaging with single-cell insulin secretion measurements will be necessary to determine how the changes in functional heterogeneity translate into hormone output. Second, the interpretation of local α-β signaling is based on spatial proximity and GLP-1R sensitivity rather than direct measurement of the secreted ligand. Finally, the mechanistic link between GLP-1R activation, cAMP signaling or other downstream events, and the observed Ca² phenotypes remain to be established experimentally.

## Conclusions

We find that intra-islet β-cell GLP-1 receptor signaling acts as a physiological amplifier and organizer of β-cell Ca^2+^ responses. It influences both the transitionary 1^st^ phase and the sustained oscillatory 2^nd^ phase of the islet’s response to glucose, more so in the α-neighboring β-cells. These effects were more apparent in pancreatic slices likely due to the preservation of α-cell content and uniform glucose exposure to all the imaged cells, suggesting that preservation of both α- and β-cells during islet isolation may improve the functional quality of islets used for transplantation. Preserving GLP-1R-mediated α-β-cell communication is important for optimal individual β-cell responsiveness, local α-cell-associated function, and multicellular β-cell network organization. Our findings provide new insight into islet physiology and the mechanisms underlying GLP-1-based therapies.

## Acknowledgements

We thank Dr. Marko Sterk of the University of Maribor, Slovenia for their involvement in developing the network analysis algorithm. We also thank Dr. Richard K.P. Benninger for his support with experimental design, reagents, and manuscript editing.

## Author Contributions

N.B. performed data analysis, collected experimental data, and wrote the manuscript; L.K.B. prepared tissue slices, collected experimental data, edited manuscript; S.B. performed glucose perfusion experiments, analyzed the data and edited the manuscript; D.R. wrote analysis code and performed data analysis; M.D. analyzed the data; M.S.K., E.P.L., J.K., J.K., M.S.H. collected experimental data, Y.Z., Y.S., A.S., A.T., D.P., A.F., S.V. performed data analysis, C.M.C. and J.D. aided in experimental design, data collection, and edited manuscript; B.P.C., R.S., A.J. maintained and provided GLP-1R KO mice; B.P.C., S.S., M.G., A.S. aided in the study design and edited the text, V.K. designed the study, wrote analysis code, collected and analyzed data, edited text. Dr. Vira Kravets is the guarantor of this work and, as such, had full access to all the data in the study and takes responsibility for the integrity of the data and the accuracy of the data analysis.

The authors N.B., L.K.B., S.B., D.R., M.D., M.S.K., E.P.L., J.K., M.S.H., Y.Z., Y.S., A.S., A.T., D.P., A.F., S.V., C.M.C., J.D., B.P.C., R.S., A.J., S.S., M.G., A.S., V.K. have no potential conflicts of interest relevant to this article to disclose.

## References

1. Benninger RKP, Kravets V: The physiological role of β-cell heterogeneity in pancreatic islet function. Nature Reviews Endocrinology 2022;18:9–22

2. Farnsworth NL, Hemmati A, Pozzoli M, Benninger RKP: Fluorescence recovery after photobleaching reveals regulation and distribution of connexin36 gap junction coupling within mouse islets of Langerhans. The Journal of Physiology 2014;592:4431–4446

3. Salomon D, Meda P: Heterogeneity and contact-dependent regulation of hormone secretion by individual B cells. Experimental Cell Research 1986;162:507–520

4. Katsuta H, Aguayo-Mazzucato C, Katsuta R, Akashi T, Hollister-Lock J, Sharma AJ, Bonner-Weir S, Weir GC: Subpopulations of GFP-Marked Mouse Pancreatic β-Cells Differ in Size, Granularity, and Insulin Secretion. Endocrinology 2012;153:5180–5187

5. Karaca M, Castel J, Tourrel-Cuzin C, Brun M, Géant A, Dubois M, Catesson S, Rodriguez M, Luquet S, Cattan P, Lockhart B, Lang J, Ktorza A, Magnan C, Kargar C: Exploring Functional β-Cell Heterogeneity In Vivo Using PSA-NCAM as a Specific Marker. PLOS ONE 2009;4:e5555

6. Kravets V, Dwulet JM, Schleicher WE, Hodson DJ, Davis AM, Pyle L, Piscopio RA, Sticco-Ivins M, Benninger RKP: Functional architecture of pancreatic islets identifies a population of first responder cells that drive the first-phase calcium response. PLoS Biol 2022;20:e3001761

7. Johnston Natalie R, Mitchell Ryan K, Haythorne E, Pessoa Maria P, Semplici F, Ferrer J, Piemonti L, Marchetti P, Bugliani M, Bosco D, Berishvili E, Duncanson P, Watkinson M, Broichhagen J, Trauner D, Rutter Guy A, Hodson David J: Beta Cell Hubs Dictate Pancreatic Islet Responses to Glucose. Cell Metabolism 2016;24:389–401

8. Salem V, Silva LD, Suba K, Georgiadou E, Neda Mousavy Gharavy S, Akhtar N, Martin-Alonso A, Gaboriau DCA, Rothery SM, Stylianides T, Carrat G, Pullen TJ, Singh SP, Hodson DJ, Leclerc I, Shapiro AMJ, Marchetti P, Briant LJB, Distaso W, …, Rutter GA: Leader β-cells coordinate Ca2+ dynamics across pancreatic islets in vivo. Nature Metabolism 2019;1:615–629

9. Bader E, Migliorini A, Gegg M, Moruzzi N, Gerdes J, Roscioni SS, Bakhti M, Brandl E, Irmler M, Beckers J, Aichler M, Feuchtinger A, Leitzinger C, Zischka H, Wang-Sattler R, Jastroch M, Tschöp M, Machicao F, Staiger H, …, Lickert H: Identification of proliferative and mature β-cells in the islets of Langerhans. Nature 2016;535:430–434

10. Benninger RKP, Hodson DJ: New Understanding of β-Cell Heterogeneity and In Situ Islet Function. Diabetes 2018;67:537–547

11. Caicedo A: Paracrine and autocrine interactions in the human islet: More than meets the eye. Seminars in Cell & Developmental Biology 2013;24:11–21

12. Huising MO, van der Meulen T, Huang JL, Pourhosseinzadeh MS, Noguchi GM: The Difference δ-Cells Make in Glucose Control. Physiology 2018;33:403–411

13. Moens K, Flamez D, Van Schravendijk C, Ling Z, Pipeleers D, Schuit F: Dual glucagon recognition by pancreatic beta-cells via glucagon and glucagon-like peptide 1 receptors. Diabetes 1998;47:66–72

14. Marchetti P, Lupi R, Bugliani M, Kirkpatrick CL, Sebastiani G, Grieco FA, Del Guerra S, D’Aleo V, Piro S, Marselli L, Boggi U, Filipponi F, Tinti L, Salvini L, Wollheim CB, Purrello F, Dotta F: A local glucagon-like peptide 1 (GLP-1) system in human pancreatic islets. Diabetologia 2012;55:3262–3272

15. Balasenthilkumaran NV, Whitesell JC, Pyle L, Friedman RS, Kravets V: Network approach reveals preferential T-cell and macrophage association with α-linked β-cells in early stage of insulitis in NOD mice. Frontiers in Network Physiology 2024;4

16. Benkahla MA, Sabouri S, Kiosses WB, Rajendran S, Quesada-Masachs E, von Herrath MG: HLA class I hyper-expression unmasks beta cells but not alpha cells to the immune system in pre-diabetes. Journal of Autoimmunity 2021;119:102628

17. Nunemaker CS, Zhang M, Wasserman DH, McGuinness OP, Powers AC, Bertram R, Sherman A, Satin LS: Individual Mice Can Be Distinguished by the Period of Their Islet Calcium Oscillations : Is There an Intrinsic Islet Period That Is Imprinted In Vivo? Diabetes 2005;54:3517–3522

18. Nunemaker CS, Wasserman DH, McGuinness OP, Sweet IR, Teague JC, Satin LS: Insulin secretion in the conscious mouse is biphasic and pulsatile. Am J Physiol Endocrinol Metab 2006;290:E523–529

19. Curry DL, L. BL, Grodsky GM: Dynamics of Insulin Secretion by the Perfused Rat Pancreas. Endocrinology 1968;83:572–584

20. Porte D, Pupo AA: Insulin responses to glucose: evidence for a two pool system in man. J Clin Invest 1969;48:2309–2319

21. Stožer A, Gosak M, Dolenšek J, Perc M, Marhl M, Rupnik MS, Korošak D: Functional connectivity in islets of Langerhans from mouse pancreas tissue slices. PLoS Comput Biol 2013;9:e1002923

22. Zmazek J, Klemen MS, Markovič R, Dolenšek J, Marhl M, Stožer A, Gosak M: Assessing Different Temporal Scales of Calcium Dynamics in Networks of Beta Cell Populations. Frontiers in Physiology 2021;12

23. Suba K, Patel Y, Martin-Alonso A, Hansen B, Xu X, Roberts A, Norton M, Chung P, Shrewsbury J, Kwok R, Kalogianni V, Chen S, Liu X, Kalyviotis K, Rutter GA, Jones B, Minnion J, Owen BM, Pantazis P, …, Salem V: Intra-islet glucagon signalling regulates beta-cell connectivity, first-phase insulin secretion and glucose homoeostasis. Mol Metab 2024;85:101947

24. Shilleh AH, Viloria K, Broichhagen J, Campbell JE, Hodson DJ: GLP1R and GIPR expression and signaling in pancreatic alpha cells, beta cells and delta cells. Peptides 2024;175:171179

25. Hart RG, Lee JJ, Zhai K, Lee S, Chauhan R, Hosseini A, Nguyen AD, Huising MO: Somatostatin receptors shape insulin and glucagon output within the pancreatic islet in mice through direct and paracrine effects. Diabetologia 2026;69:2569–2584

26. Denwood G, Tarasov A, Salehi A, Vergari E, Ramracheya R, Takahashi H, Nikolaev VO, Seino S, Gribble F, Reimann F, Rorsman P, Zhang Q: Glucose stimulates somatostatin secretion in pancreatic δ-cells by cAMP-dependent intracellular Ca2+ release. Journal of General Physiology 2019;151

27. Stožer A, Dolenšek J, Rupnik MS: Glucose-Stimulated Calcium Dynamics in Islets of Langerhans in Acute Mouse Pancreas Tissue Slices. PLOS ONE 2013;8:e54638

28. Saikia M, Holter MM, Donahue LR, Lee IS, Zheng QC, Wise JL, Todero JE, Phuong DJ, Garibay D, Coch R, Sloop KW, Garcia-Ocana A, Danko CG, Cummings BP: GLP-1 receptor signaling increases PCSK1 and β cell features in human α cells. JCI Insight 2021;6:141851

29. Schindelin J, Arganda-Carreras I, Frise E, Kaynig V, Longair M, Pietzsch T, Preibisch S, Rueden C, Saalfeld S, Schmid B, Tinevez J-Y, White DJ, Hartenstein V, Eliceiri K, Tomancak P, Cardona A: Fiji: an open-source platform for biological-image analysis. Nature Methods 2012;9:676–682

30. Karlöf L, Ølgård TA, Godtliebsen F, Kaczmarska M, Fischer H: Statistical techniques to select detection thresholds for peak signals in ice-core data. Journal of Glaciology 2005;51:655–662

31. Šterk M, Zhang Y, Pohorec V, Leitgeb EP, Dolenšek J, Benninger RKP, Stožer A, Kravets V, Gosak M: Network representation of multicellular activity in pancreatic islets: Technical considerations for functional connectivity analysis. PLOS Computational Biology 2024;20:e1012130

32. Šterk M, Dolenšek J, Skelin Klemen M, Križančić Bombek L, Paradiž Leitgeb E, Kerčmar J, Perc M, Slak Rupnik M, Stožer A, Gosak M: Functional characteristics of hub and wave-initiator cells in β cell networks. Biophys J 2023;122:784–801

33. Zhang Q, Galvanovskis J, Abdulkader F, Partridge CJ, Göpel SO, Eliasson L, Rorsman P: Cell coupling in mouse pancreatic beta-cells measured in intact islets of Langerhans. Philos Trans A Math Phys Eng Sci 2008;366:3503–3523

34. Duh M, Šterk M, Križančić Bombek L, MacDonald PE, Stožer A, Gosak M: Spatially bound functional heterogeneity drives modular organization in β-cell networks. Biophysical Journal 2025;124:3008–3022

35. Jaffredo M, Bertin E, Pirog A, Puginier E, Gaitan J, Oucherif S, Lebreton F, Bosco D, Catargi B, Cattaert D, Renaud S, Lang J, Raoux M: Dynamic uni- and multicellular patterns encode biphasic activity in pancreatic islets. Diabetes 2021;70:878–888

36. Ren H, Li Y, Han C, Yu Y, Shi B, Peng X, Zhang T, Wu S, Yang X, Kim S, Chen L, Tang C: Pancreatic α and β cells are globally phase-locked. Nat Commun 2022;13:3721

37. Briggs JK, Gresch A, Marinelli I, Dwulet JM, Albers DJ, Kravets V, Benninger RKP: β- cell intrinsic dynamics rather than gap junction structure dictates subpopulations in the islet functional network. Elife 2023;12

38. Cerasi E: POTENTIATION OF INSULIN RELEASE BY GLUCOSE IN MAN: I. Quantitative analysis of the enhancement of glucose-induced insulin secretion by pretreatment with glucose in normal subjects. Acta Endocrinologica 1975;79:483–501

39. Niki I, Tamagawa T, Niki H, Niki A, Koide T, Sakamoto N: Possible involvement of diacylglycerol-activated, Ca2+-dependent protein kinase in glucose memory of the rat pancreatic B-cell. Acta Endocrinologica 1988;118:204–208

40. Grill V, Rundfeldt M: Effects of Priming with D-Glucose on Insulin Secretion from Rat Pancreatic Islets: Increased Responsiveness to Other Secretagogues. Endocrinology 1979;105:980–987

41. Santos GJd, Ferreira SM, Ortis F, Rezende LF, Li C, Naji A, Carneiro EM, Kaestner KH, Boschero AC: Metabolic memory of ß-cells controls insulin secretion and is mediated by CaMKIIa. Molecular Metabolism 2014;3:484–489

42. Tian G, Sandler S, Gylfe E, Tengholm A: Glucose- and hormone-induced cAMP oscillations in α- and β-cells within intact pancreatic islets. Diabetes 2011;60:1535–1543

43. Stožer A, Paradiž Leitgeb E, Pohorec V, Dolenšek J, Križančić Bombek L, Gosak M, Skelin Klemen M: The Role of cAMP in Beta Cell Stimulus-Secretion and Intercellular Coupling. Cells 2021;10

44. Wan QF, Dong Y, Yang H, Lou X, Ding J, Xu T: Protein kinase activation increases insulin secretion by sensitizing the secretory machinery to Ca2+. J Gen Physiol 2004;124:653–662

45. Kasai H, Hatakeyama H, Ohno M, Takahashi N: Exocytosis in islet beta-cells. Adv Exp Med Biol 2010;654:305–338

46. Light PE, Manning Fox JE, Riedel MJ, Wheeler MB: Glucagon-like peptide-1 inhibits pancreatic ATP-sensitive potassium channels via a protein kinase A- and ADP-dependent mechanism. Mol Endocrinol 2002;16:2135–2144

47. Shuai H, Xu Y, Ahooghalandari P, Tengholm A: Glucose-induced cAMP elevation in β-cells involves amplification of constitutive and glucagon-activated GLP-1 receptor signalling. Acta Physiol (Oxf) 2021;231:e13611

48. Capozzi ME, Svendsen B, Encisco SE, Lewandowski SL, Martin MD, Lin H, Jaffe JL, Coch RW, Haldeman JM, MacDonald PE, Merrins MJ, D’Alessio DA, Campbell JE: β Cell tone is defined by proglucagon peptides through cAMP signaling. JCI Insight 2019;4

49. Svendsen B, Larsen O, Gabe MBN, Christiansen CB, Rosenkilde MM, Drucker DJ, Holst JJ: Insulin Secretion Depends on Intra-islet Glucagon Signaling. Cell Rep 2018;25:1127–1134.e1122

50. Capozzi ME, Wait JB, Koech J, Gordon AN, Coch RW, Svendsen B, Finan B, D’Alessio DA, Campbell JE: Glucagon lowers glycemia when β-cells are active. JCI Insight 2019;5

