## Supplemental Figure for "Role of GLP1-receptor signaling and α-β communication in functional β-cell heterogeneity"

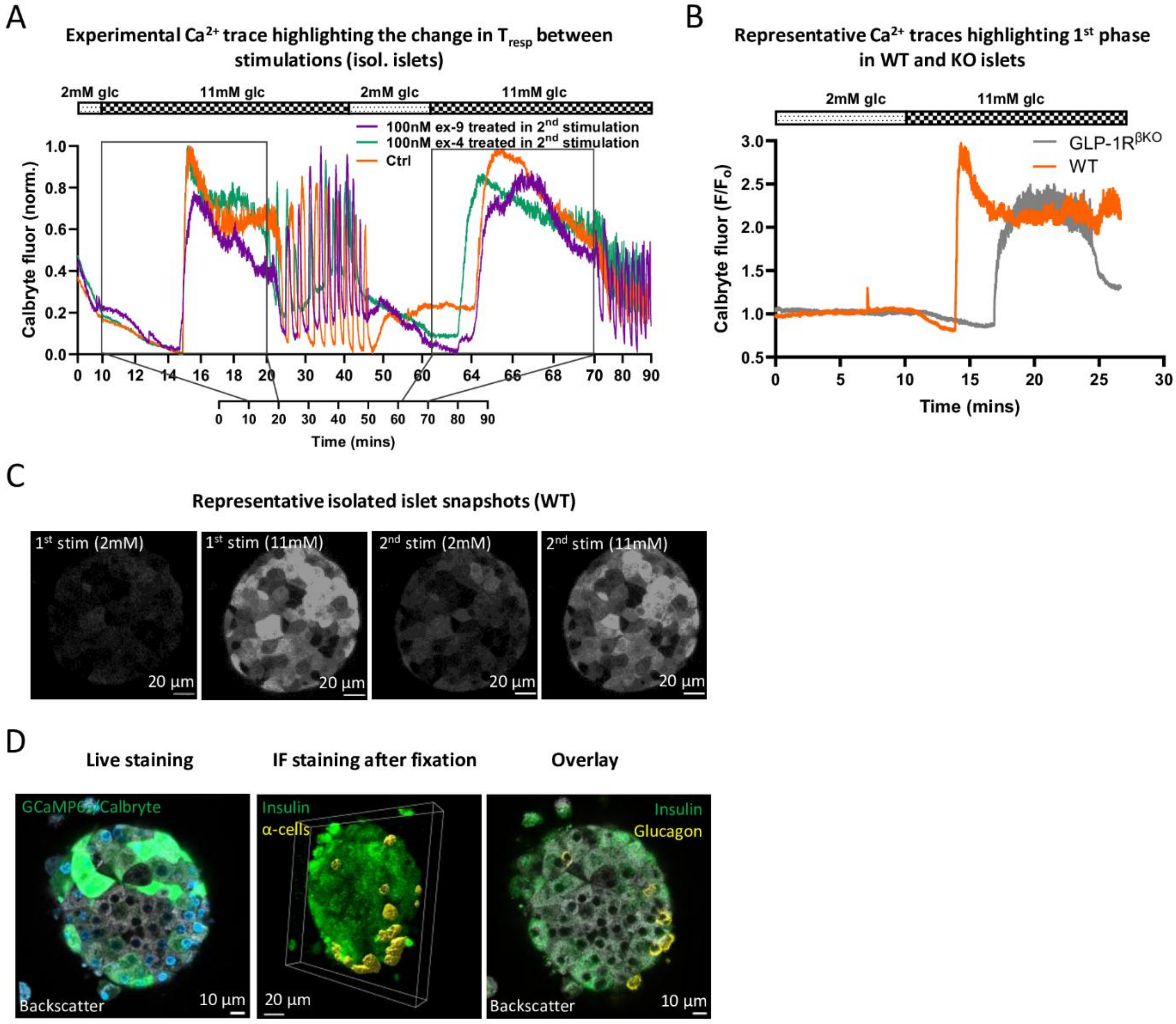


**Supplementary Figure. S1 (A)** Experimental protocol for the different treatments with representative islet-average [$Ca^{2+}$] time courses highlighting the 1st phase response in isolated islets. Traces were aligned to the onset of the 1st glucose-induced 1st phase response to facilitate visual comparison of the 2nd stimulation. **(B)** Representative islet average $[Ca]^{2+}$ traces from WT and KO islets highlighting the 1st phase. **(C)** Representative islet snapshots at selected time points from GCaMP6s isolated islets. **(D)** Representative GCAMP6s isolated islet used for [$Ca^{2+}$] time-series imaging; 3D rendering of the IF z-stack after fixation; live and fixed image overlays.

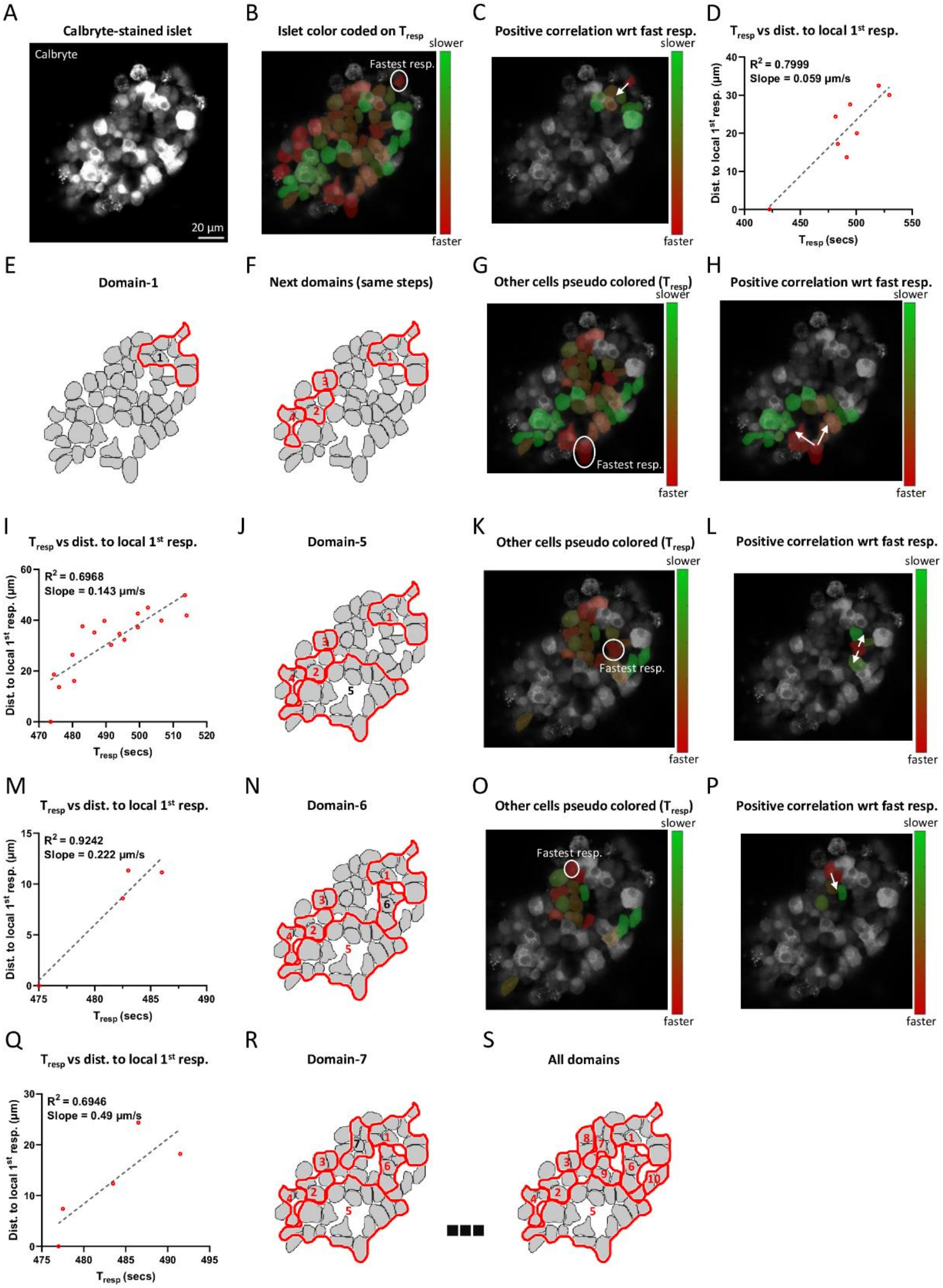


**Supplementary Figure. S2.** Schematic representation of the bias-free domain creation algorithm.

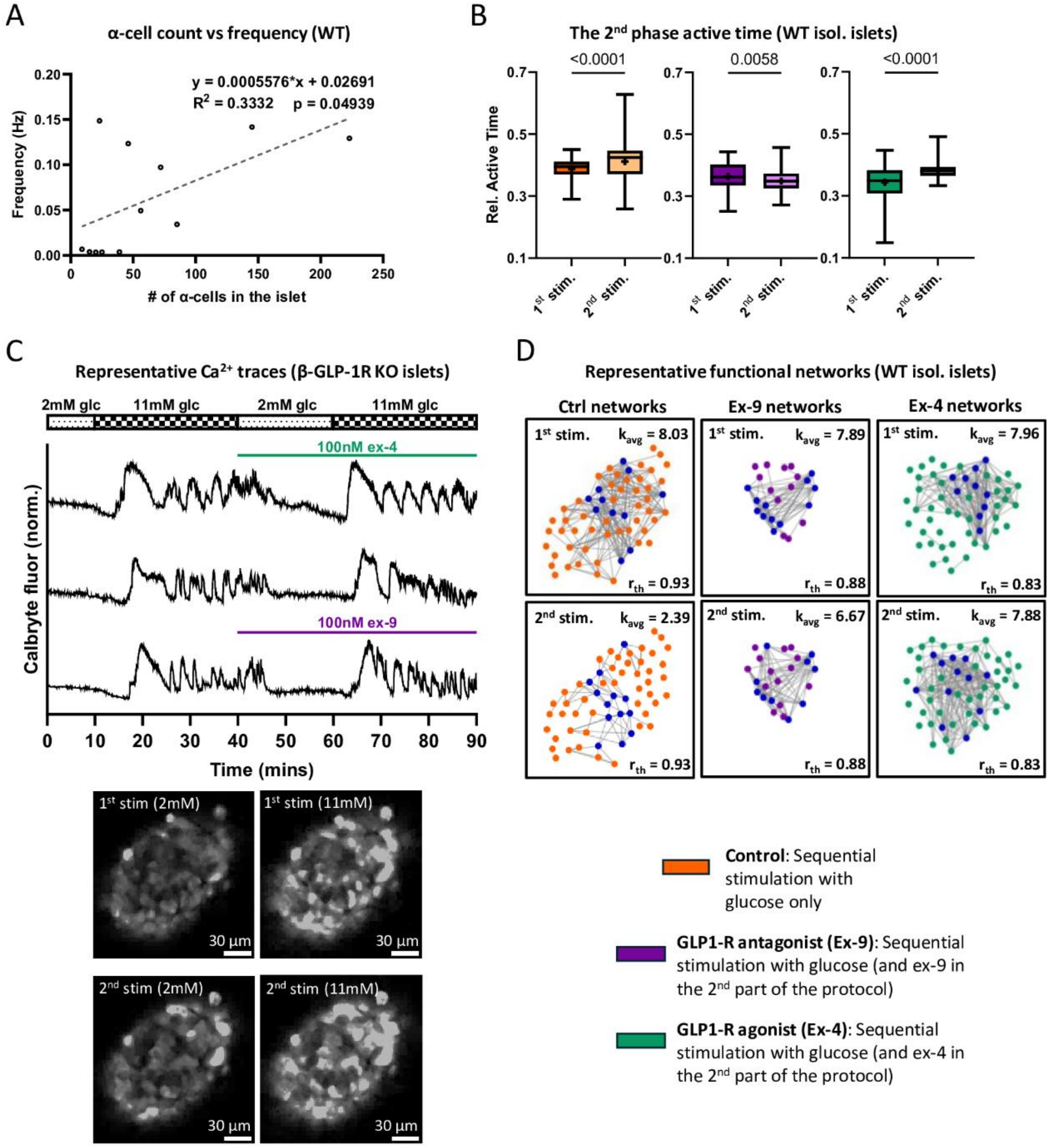


**Supplementary Figure. S3 (A)** Relationship between the islet-average frequency of [$Ca^{2+}$] oscillations and number of α-cells in the islet (n = 12 islets, n = 5 from isolated islets and n = 7 from pancreatic slices. Simple linear regression was used for statistical analysis. **(B)** Relative active time of each β-cell in 1st vs 2nd stimulations in control, ex-9-treated, and ex-4-treated islets (n = 5 control, n = 5 ex-9, and n = 5 ex-4 treated islets). Wilcoxon matched-pairs signed-rank test was used for statistical analysis. **(C)** Experimental protocol for the different treatments, with islet-average [$Ca^{2+}$] time courses and representative islet snapshots of KO islets at selected time points. **(D)** Functional β-cell networks in the 1st and 2nd stimulations for representative control, ex-9 treated and ex-4 treated isolated islets.
